# Global emergence and population dynamics of divergent serotype 3 CC180 pneumococci

**DOI:** 10.1101/314880

**Authors:** Taj Azarian, Patrick K Mitchell, Maria Georgieva, Claudette M Thompson, Amel Ghouila, Andrew J Pollard, Anna von Gottberg, Mignon du Plessis, Martin Antonio, Brenda A Kwambana-Adams, Stuart C Clarke, Dean Everett, Jennifer Cornick, Ewa Sadowy, Waleria Hryniewicz, Anna Skoczynska, Jennifer C Moïsi, Lesley McGee, Bernard Beall, Benjamin J Metcalf, Robert F Breiman, PL Ho, Raymond Reid, Kate L O’Brien, Rebecca A Gladstone, Stephen D Bentley, William P Hanage

**Affiliations:** Center for Communicable Disease Dynamics, Department of Epidemiology, T.H. Chan School of Public Health, Harvard University, Boston, MA; Institut Pasteur de Tunis, LR11IPT02, Laboratory of Transmission, Control and Immunobiology of Infections (LTCII), Tunis-Belvédère, Tunisia; Oxford Vaccine Group, Department of Paediatrics, University of Oxford; NIHR Oxford Biomedical Research Centre, Centre for Clinical Vaccinology and Tropical Medicine (CCVTM), Churchill Hospital, Oxford OX3 7LJ, UK; Centre for Respiratory Diseases and Meningitis, National Institute for Communicable Diseases of the National Health Laboratory Service, Johannesburg, South Africa; Medical Research Council Unit The Gambia, Atlantic Road, Fajara, The Gambia; Faculty of Medicine and Institute for Life Sciences and Global Health Research Institute, University of Southampton, S016 6YD, United Kingdom; NIHR Southampton Biomedical Research Centre; Queens Research Institute, University of Edinburgh; Institute of Infection and Global Health, University of Liverpool, Liverpool, UK; National Medicines Institute, Warsaw Poland; Agence de Médecine Préventive, Paris, France; Respiratory Diseases Branch, Centers for Disease Control and Prevention, Atlanta, GA 30333, USA; Global Health Institute, Emory University, Atlanta GA; Department of Microbiology, Queen Mary Hospital University of Hong Kong, Hong Kong, People’s Republic of China; Center for American Indian Health, Johns Hopkins Bloomberg School of Public Health, Baltimore, Maryland; The Wellcome Trust Sanger Institute, Wellcome Trust Genome Campus, Hinxton, Cambridge CB10 1SA, UK

## Abstract

*Streptococcus pneumoniae* serotype 3 remains a significant cause of morbidity and mortality worldwide, despite inclusion in the 13-valent pneumococcal conjugate vaccine (PCV13). Serotype 3 increased in carriage since the implementation of PCV13 in the United States, while invasive disease rates remain unchanged. We investigated the persistence of serotype 3 in carriage and disease, through genomic analyses of a global sample of 301 serotype 3 isolates of the Netherlands^3^–31 (PMEN31) clone CC180, combined with associated patient data and PCV utilization among countries of isolate collection. We assessed phenotypic variation between dominant clades in capsule charge (zeta potential), capsular polysaccharide shedding, and susceptibility to opsonophagocytic killing, which have previously been associated with carriage duration, invasiveness, and vaccine escape. We identify a recent shift in the CC180 population attributed to a lineage termed Clade II, which was estimated by Bayesian coalescent analysis to have first appeared in 1968 [95% HPD: 1939–1989] and increased in prevalence and effective population size thereafter. Clade II isolates are divergent from the pre-PCV13 serotype 3 population in non-capsular antigenic composition, competence, and antibiotic susceptibility, the last resulting from the acquisition of a Tn*916*-like conjugative transposon. Differences in recombination rates among clades correlated with variations in the ATP-binding subunit of Clp protease as well as amino acid substitutions in the *comCDE* operon. Opsonophagocytic killing assays elucidated the low observed efficacy of PCV13 against serotype 3. Variation in PCV13 use among sampled countries was not independently correlated with the CC180 population shift; therefore, genotypic and phenotypic differences in protein antigens and, in particular, antibiotic resistance may have contributed to the increase of Clade II. Our analysis emphasizes the need for routine, representative sampling of isolates from disperse geographic regions, including historically under-sampled areas. We also highlight the value of genomics in resolving antigenic and epidemiological variations within a serotype, which may have implications for future vaccine development.

**Author Summary:** *Streptococcus pneumoniae* is a leading cause of bacterial pneumoniae, meningitis, and otitis media. Despite inclusion in the most recent pneumococcal conjugate vaccine, PCV13, serotype 3 remains epidemiologically important globally. We investigated the persistence of serotype 3 using whole-genome sequencing data form 301 isolates collected among 24 countries from 1993–2014. Through phylogenetic analysis, we identified three distinct lineages within a single clonal complex, CC180, and found one has recently emerged and grown in prevalence. We then compared genomic difference among lineages as well as variations in pneumococcal vaccine use among sampled countries. We found that the recently emerged lineage, termed Clade II, has a higher prevalence of antibiotic resistance compared to other lineages, diverse surface protein antigens, and a higher rate of recombination, a process by which bacteria can uptake and incorporate genetic material from its surroundings. Differences in vaccine use among sampled countries did not appear to be associated with the emergence of Clade II. We highlight the need to routine, representative sampling of bacterial isolates from diverse geographic areas and show the utility of genomic data in resolving epidemiological differences within a pathogen population.

## Introduction

Pneumococcal disease caused by *Streptococcus pneumoniae* remains a significant cause of morbidity and mortality even in the era of effective conjugate vaccines, which protect against up to 13 different serotypes. Among the serotypes covered by the 13-valent pneumococcal conjugate vaccine (PCV13), serotype 3 is considered highly invasive and is associated with a high risk of mortality in both observational studies and animal models [1–4]. However, the serotype-specific effectiveness of PCV13 against serotype 3 remains uncertain. There has been little reduction in serotype 3 disease compared to disease due to other vaccine serotypes following implementation of PCV13 in the US (Figure 1) [5]. In addition, data from a randomized controlled trials on nasopharyngeal carriage [6], post-licensure studies of PCV13 on invasive pneumococcal disease (IPD) [7], and national surveillance data on IPD from England and Wales [8,9], among other studies have shown little vaccine effect on serotype 3.

**Figure 1.**
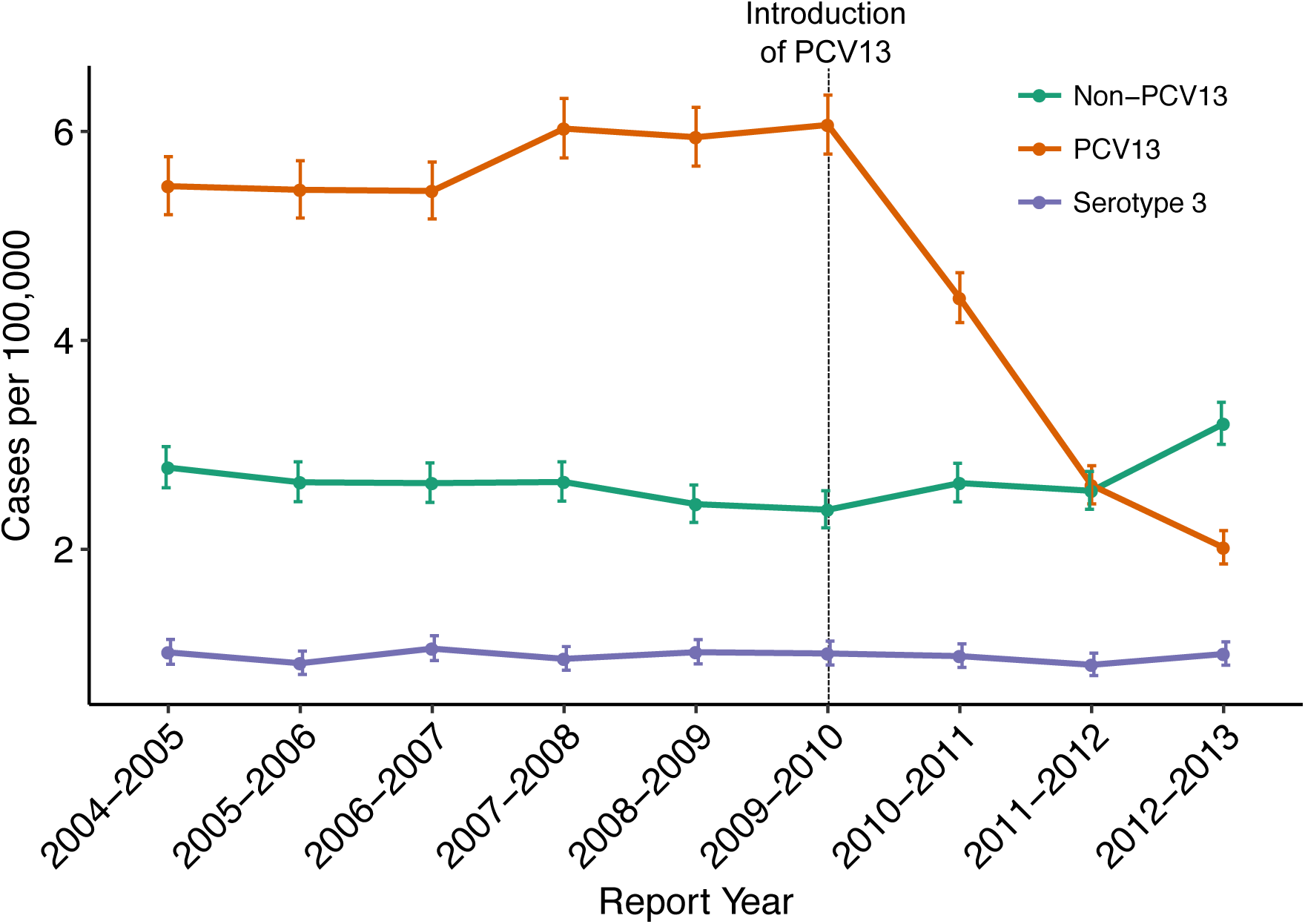
Changes in the incidence of invasive pneumococcal disease (IPD) from 2004 through 2013 among all ages and vaccine status in the United States based on CDC Active Bacterial Core (ABC) surveillance data. [5]. Rates of IPD expressed as cases per 100,000 population are on the y-axis, and calendar year of surveillance on the x-axis. Green line represents the IPD rate for the five most common non-PCV13 serotypes; purple, the rate for serotype 3 only; and orange, the rate of (PCV13 – serotype 3) serotypes containing (1, 4, 5, 6A, 6B, 7 F, 9 V, 14, 18 C, 19A, 19 F, and 23 F). Further information can be obtained from CDC’s ABC surveillance (www.cdc.gov/abcs/).

Multi-locus sequence typing (MLST) data has shown that the majority of serotype 3 isolates globally are a single clonal complex (CC) of closely related genotypes (CC180) also known as the Netherlands^3^–31 (PMEN31) clone [10–13], with the exception of Africa where non-CC180 MLST types are prevalent [14]. However, more recent genomic studies have shown that CC180 contains multiple distinct lineages [12,15], which MLST is not sufficiently discriminating to detect. While one of these lineages (termed ‘Clade I’) accounts for the overwhelming majority of previously studied genomes, the overwhelming majority (92%) of these isolates came from European collections, and might not be representative of the global pneumococcal population.

Following roll-out of PCV13, an unexpected increase in the carriage prevalence of serotype 3 was noted among Massachusetts children less than seven years of age [16]. Initial genomic analysis of these carriage isolates showed a change in the serotype 3 population [17]. Prior to vaccination the majority of isolates fell into the previously described ‘Clade I’, while following PCV13 introduction, the observed increase in carriage prevalence was largely due to isolates which, while still CC180, were drawn from a more diverse population that has been poorly represented in prior samples. Phenotypic variations within CC180 and how they may relate to vaccine efficacy have not been investigated and remain poorly understood.

To investigate the population structure and evolutionary history of serotype 3 CC180 pneumococci, we conducted a genomic analysis of 301 serotype 3 CC180 isolates from carriage and disease collected from 24 countries over 20 years. Further, we assessed multiple phenotypic features linked to the epidemiology of serotype 3, and which might contribute to vaccine escape including reaction to PCV13 antisera, and how they vary in the CC180 population [18–20].

## Methods

### Study Population and Epidemiological data

The sample consisted of 301 CC180 (PMEN31) genomes collected from carriage (n=68), invasive disease (n=231), and unknown clinical manifestations (n=2) between 1993–2014. Whole-genome sequencing (WGS) data for this sample were obtained from a previous analysis of serotype 3 described elsewhere (n=82) [12], on-going studies of carriage in Massachusetts, USA (n=27) [21], carriage studies in the South West United States (n=14) [22], the Bacterial Isolate Genome Sequence database (BIGSDB) (n=10), and the Global Pneumococcal Sequencing (GPS) project (http://www.pneumogen.net/gps/) (n=168). Available data included the year and country of isolation, isolation source, and limited patient demographic data. To assess the temporal shift in prevalence of the three CC180 clades, we tested the significance of changes in the proportion of isolates by clade for each sampling year by comparing yearly proportions to 1000 random deviates of a Dirichlet distribution [23]. If the temporal change in proportion was found to be significant, then the directionality of the change was assessed. Current and past PCV (PCV7, PCV10, and PCV13) use data for countries where the serotype 3 CC180 sample was collected were queried from International Vaccine Access Center (IVAC) VIEW-hub website (www.view-hub.org, accessed April 5, 2017). Fisher’s exact test was used to test the association between countries that introduced PCV and serotype 3 clade emergence based on phylogenetic demography (See phylogenetic analysis methods). Further, among countries that introduced PCV13, the correlation between the date of introduction and changes in serotype 3 demography was assessed using Pearson’s correlation coefficient. All statistical analysis and figure generation was performed using Rstudio v1.0.143 with R v3.3.1.

### Genomic Analysis

For GPS isolates, raw sequencing data and *de novo* assemblies were downloaded from http://www.pneumogen.net/gps/. GPS *de novo* assemblies were generated using a Velvet pipeline as previously described [24]. The remaining sequencing data were downloaded from the NCBI SRA database (See supplementary table for a list of accession numbers). If only draft assemblies were available, paired-end 150 bp reads were generated using the BBMap’s RandomReads script. *De novo* assemblies for non-GPS isolates were generated using SPAdes v3.10 as previously described [12,25]. Assemblies were then annotated using Prokka v1.12 [26] and pangenome analysis was conducted using Roary [27]. Assemblies constructed using simulated reads were checked for variation in assembly metrics, annotation, and SNP diversity, in order to detect any evidence of sequencing and assembly errors. Variants for 13 polymorphic pneumococcal protein antigens, including pneumococcal surface proteins C and A (*pspC* and *pspA*) were determined by mapping raw reads to an antigen variant database using SRST2 as previously described [22]. Quality filtered and trimmed reads were mapped to *S. pneumoniae* OXC141 (NCBI Reference Sequence: NC_017592), a Clade I, serotype 3 ST180/CC180 carriage isolate, using SMALT v0.7.6. SNPs were called using SAMtools v1.3.1 [28], and were filtered requiring QUAL>50, depth of coverage ≥5 and a minimum alternate allele frequency ≥0.75 [21,29]. Gubbins v2.1 was used to assess recombination, and coding sequences (CDSs) impacted by recombination blocks were annotated and plotted using Circos [30,31]. Last, we assessed non-synonymous mutations and recombination in the *comCDE* operon, which has previously been implicated in the low competence of CC180 compared to other pneumococcal lineages [12].

### Phylogenetic Analysis

A maximum likelihood (ML) phylogeny was inferred from a recombination-censored SNP alignment using RAxML v8.2.1 with an ASC_GTRGAMMA nucleotide substitution model, Lewis ascertainment bias correction, and 100 bootstrap replicates [32]. The ML phylogeny was rooted using strain AP200 (accession # CP002121), a serotype 11A, ST62 *S. pneumoniae* invasive isolate, which was found to be immediately basal to the CC180 clade [33]. After assigning isolates to major clades, Gubbins was run independently on the sequence alignments from each clade to identify putative recombination events. ML phylogenies from recombination-censored alignments were used to test temporal signal by assessing correlation between strain isolation date and root-to-tip distance. To reduce bias in coalescent analysis due to differences in sampling over the study period, isolates were subsampled using a uniform probability from each year and spatial location (country and region), as recommended in [34]. To verify temporal signal, we performed date randomization tests on the down-sampled datasets, whereby the sampling times of sequences are shuffled and evolutionary rates are compared to correctly assigned time estimates (see supplementary methods) [35,36]. For major Clade I (hereon referred to as Clade I-α) and Clade II, coalescent analysis was performed using BEAST v1.8.4 (see supplemental methods) [37]. Parameter estimates for the evolutionary rate, root height, and *N_e_* were obtained from the best-fit model and compared between clades. Last, to infer the ancestral geographic location and migration history of Clade II, we used the structured coalescent implemented in Beast 2.4.4 specifying the region of collection as the geographic location for each tip [38,39]. Additionally, BEAST XML code has been made available (https://github.com/c2-d2/Projects).

### Phenotypic and Genotypic Antibiotic Resistance

Antimicrobial resistance (AMR)-associated genes and SNPs were identified with ARIBA using ARG-ANNOT and CARD databases [40,41]. Penicillin MICs were predicted using WGS data to type transpeptidase domains of penicillin binding proteins [42]. Genotypic antibiotic resistance was validated among strains with available broth dilution data using published CLSI breakpoints for penicillin, chloramphenicol, erythromycin, clindamycin, and tetracycline. For isolates that possessed multiple AMR-associated genes, annotated *de novo* assemblies were investigated to identify conjugative transposons. Transposons were identified through review of *de novo* assembly annotations and their presence/absence among strains was confirmed by mapping sequencing reads to transposons as described above.

### Assessment of Capsular Polysaccharide Variations

First, we assessed variation in the CPS loci by abstracting the region from reference-based genome assemblies and generating a ML phylogeny using RAxML with GTRGAMMA substitution model and 100 bootstrap replicates. The mean pairwise nucleotide difference was also calculated enforcing pairwise deletion of missing sites. CPS loci were manually inspected for recombination events using the output from Gubbins to determine the potential role in clade variation.

To investigate phenotypic variations related to the serotype 3 CC180 capsular polysaccharide between Clades I-α and II and potentially explain the recent emergence of Clade II, we assessed surface charge (zeta potential), capsular release, and opsonophagocytic killing. Zeta potentials among representative Clade I-α (n=5) and Clade II (n=3) serotype 3 strains from Massachusetts were compared to *S. pneumoniae* serotype 3 ST378 laboratory strain WU2 and ΔCPS WU2 as previously described [19]. Capsular release, a mechanism by which type 3 CPS interferes with antibody-mediated killing and gains protection by anti-CPS antibodies, was compared among Clade I-α (n=3) and Clade II (n=3) serotype 3 isolates from Massachusetts to strain WU2 as previously described [43]. To assess the efficiency of antisera against serotype 3 CC180 to opsonize pneumococci for uptake and killing by differentiated polymorphonuclear leukocytes, we used an opsonophagocytic killing assay (OPKA) [44]. The killing assays were performed at multiple dilutions using PCV13 antisera and antisera generated from type 3 polysaccharide (PS) (see supplemental methods) [45]. All Clade I-α and II strains studied in phenotypic assays are marked on the ML phylogeny.

## Results

### Population structure

The serotype 3 CC180 isolates included in this sample came from 24 countries from North America (38.9%), Western Europe (17.9%), Asia (14.6%), Eastern Europe (14.3%), Africa (7.6%), and South America (6.3%) collected from 1993–2014. Our phylogenetic analysis improved the resolution of the three major lineages identified in previous work. Clade I and the previously described Clade II together form a single monophyletic lineage, distinct from the previously described Clade III (Figure 2A). Applying an out-group roots the phylogeny roots on the branch between those strains previously described as clade II and III, indicating that the previous clade naming scheme was inaccurate. To reflect this, and the fact that we now find two rather than three monophyletic lineages, we refer to Clades I-α and I-β (formerly Clades I and II) and Clade II (formerly Clade III). Clade I-α isolates made up the majority of the sample (68.4%), but a significant proportion is made up of Clade I-β (12.3%) and Clade II (19.3%). The expanded sample now shows a single deep branching lineage containing Clade I-β, which is polyphyletic and subtends Clade I-α. Clade II is further subdivided into three well-supported subclades that are distinct in terms of genome content (see below). Following removal of regions that were inferred to have been introduced by recombination, the nucleotide diversity of Clade II was significantly greater than that of Clade I-α [mean pairwise SNP distance 98.0 (SE 2.1) vs. 112.7 (SE 3.6)].

**Figure 2.**
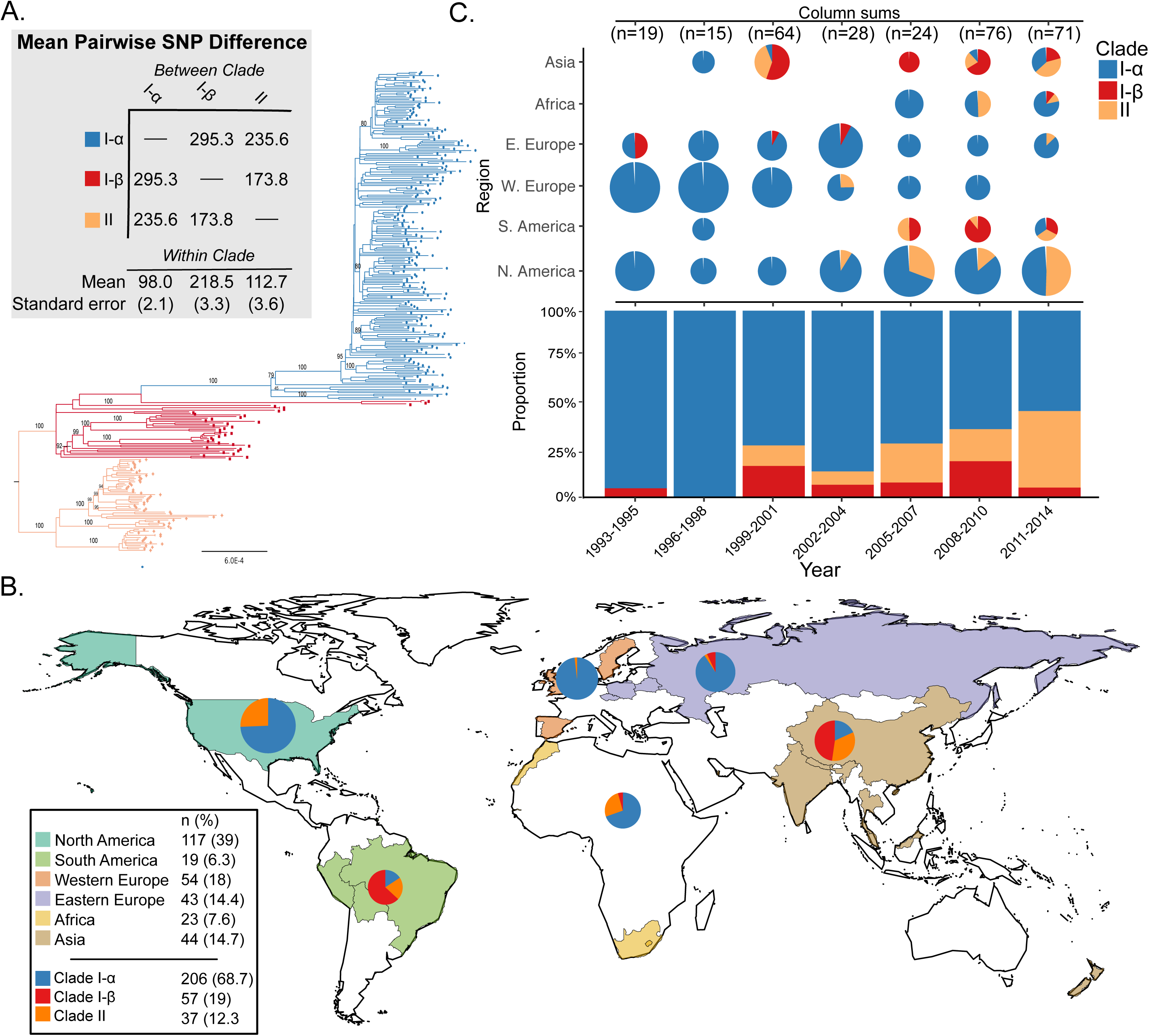
A) Rooted maximum likelihood phylogeny of *S. pneumoniae* serotype 3 CC180 isolates (n=301). Phylogeny was out-group rooted using strain AP200 (accession # CP002121) a serotype 11A, ST62 *S. pneumoniae* invasive isolate, which was found to be immediately basal to the CC180 clade in a global phylogeny of pneumococcal reference genomes and therefore represented the closest out-group. Bootstrap values from 100 replicates are labelled on major clades and epidemiologically relevant sub-clades. Additional values can be found in the supplemental phylogeny. Mean pairwise within- and between-clade SNP difference are presented in the grey shaded box with color corresponding to the phylogeny. B.) World map illustrating sampled countries and regions with respective proportion of isolates belonging to Clade I-α, I-β, and II. Countries are colored according to region and pie charts represent the proportion of isolates belonging to major serotype 3 clades. The size of the pie chart is scaled to the proportion of strains sampled from each region. C) Proportion of clade membership by three-year collection window. The proportion of clade membership by region over-time is displayed on the top of the figure. Pie charts are scaled by the number of isolates sampled from a geographic region by time window (i.e., column). The overall proportion of clades for each time window is presented on the bottom of the figure.

### Phylogeography and country-level vaccine use history

The proportion of isolates belonging to Clade II has significantly increased over time from 1999–2001 (11%) to 2011–2014 (41%) (4.2, 95% CI: 1.9 – 9.0, p<0.0001) with the largest increase occurring in North America (Figure 2B). During the same time Clade I- α has significantly decreased, and Clade I-β remained largely unchanged. In the present sample, Clade II was first reported in Asia in 1999, but is now globally distributed, making up a large proportion of samples from Asia, Africa, and North and South America (Figure 2B). However, Clade II was only observed twice among 97 European strains (Table 1), first appearing in 2003. Clade I-α was the most prevalent in the sample and found to be globally distributed, while Clade I-β isolates appear more common in samples from South America and Asia (Figure 2B).

**Table 1.** Clade composition of CC180 serotype 3 isolates by geographic location. Percentages are based on 295 isolates for which the collection date and country are known.

| Clade | Eastern Europe (n=43) | Western Europe (n=54) | North America (n=117) | South America (n=15) | Africa (n=23) | Asia (n=43) | Total |
| --- | --- | --- | --- | --- | --- | --- | --- |
| Clade I-α (n=204) | 90.7% | 98.1% | 74.4% | 6.7% | 69.6% | 18.6% | 69.2% |
| Clade I-β (n=35) | 7.0% | - | - | 73.3% | 4.3% | 46.5% | 11.9% |
| Clade II (n=56) | 2.3% | 1.9% | 25.6% | 20.0% | 26.1% | 34.9% | 19.0% |

Clades I-α and II had significant temporal signal, identified by root-to-tip date correlation (Supplemental Figures 1 and 2). A date randomization test also showed significant temporal signal for Clade II, but date randomizations for Clade I-α failed to reach ESS values >200 despite chain length (Supplemental Figures 3). Model comparison using Bayes factors calculated from MLEs identified both Clades I-α and II fit a GMRF SkyGrid demographic model and relaxed molecular clock (Supplemental Table 1). In addition, the exponential demographic model was preferred to the constant for Clade II but rejected Clade I-α. Evolutionary rates were not significantly different between the two clades; however, the 95% Highest Posterior Density (HPD) for Clade II was wider (Supplemental Figure 4), owing to a greater coefficient of variation (*S*=0.38 vs 0.21 for Clades II and I-α, respectively) and indicating higher rate variation among branches. Clade II was significantly younger, with an estimated most recent common ancestor (TMRCA) of 1968 [95% HPD: 1939–1989] (Supplemental Table 1). The effective population size (*N_e_*) for Clades I-α and II have been increasing, as demonstrated by the SkyGrid *N_e_* plots (Figure 3) and rejection of the constant population size demographic model (Supplemental Table 1). Further, the exponential model for Clade II suggested the population was exponentially increasing (mean exponential growth rate = 0.054 [95% HPD: 5.11×10^-3^, 0.11].

**Figure 3.**
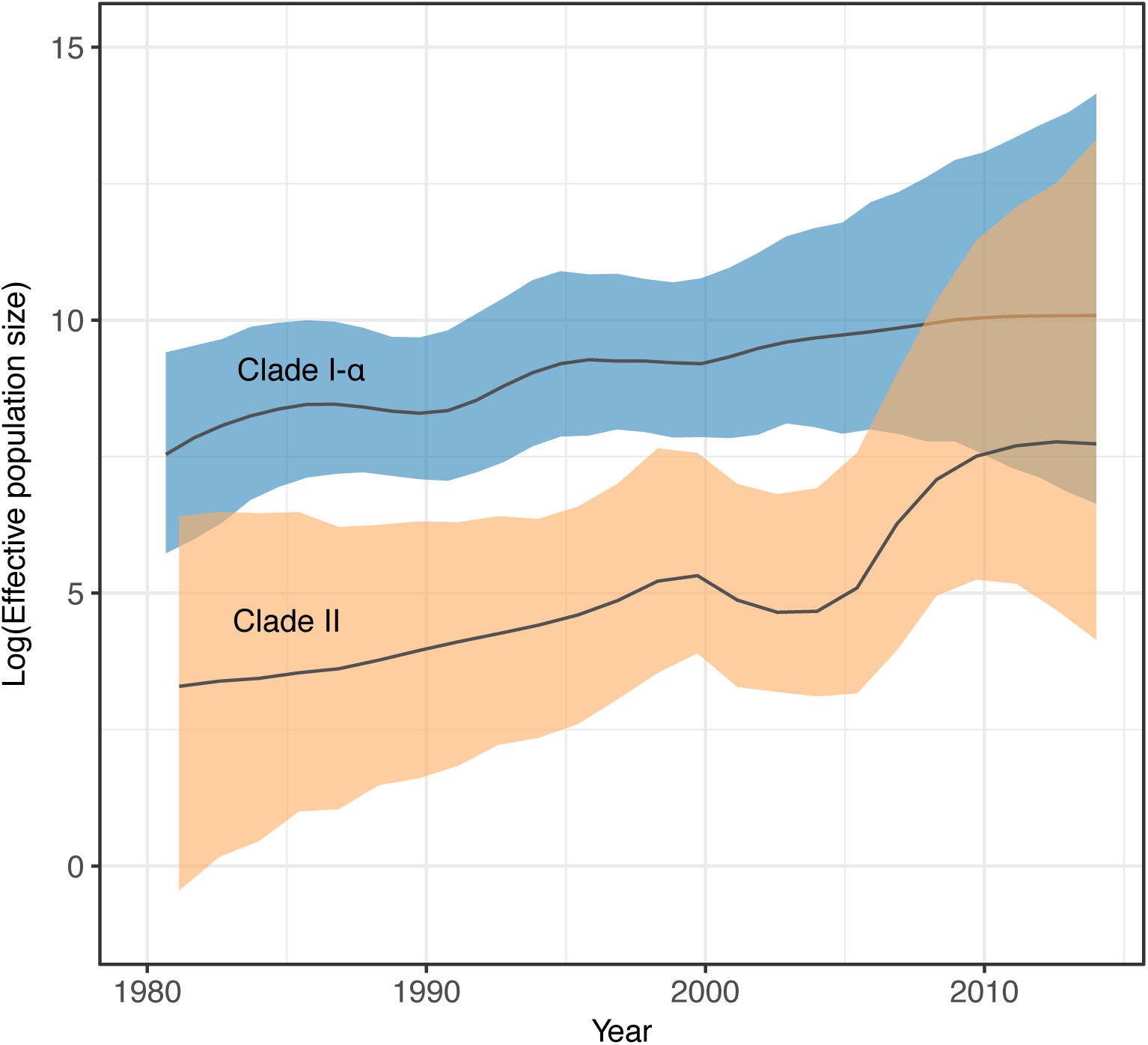
Effective population size (*N_e_*) comparison of serotype 3 CC180 Clades I-α and II. *N_e_* values were estimated using BEAST 1.8.4, enforcing a GMRF SkyGrid demographic model and relaxed molecular clock. The *N_e_* of Clades I-α and II have been increasing with Clade II in particular increasing exponentially base on phylodynamic model comparison.

Phylogeographic migration models of Clade II using the structured coalescent achieved sufficient mixing (i.e., ESS values >200) for all parameters; however, posterior probabilities for migration rates between geographic regions were not significant (<0.30). Therefore, we were unable to infer the ancestral geographic locations of Clade II isolates. We noted, however, that in both ML and Bayesian time-scaled phylogenies, the Asian clade made up of isolates from Hong Kong is proximally basal to the major North American clade (Figures 4 and 5). Further, isolates from Hong Kong possess a unique Tn*916*-like transposon shared only with isolates found in the dominant North American clade (Figure 5) (See below).

**Figure 4.**
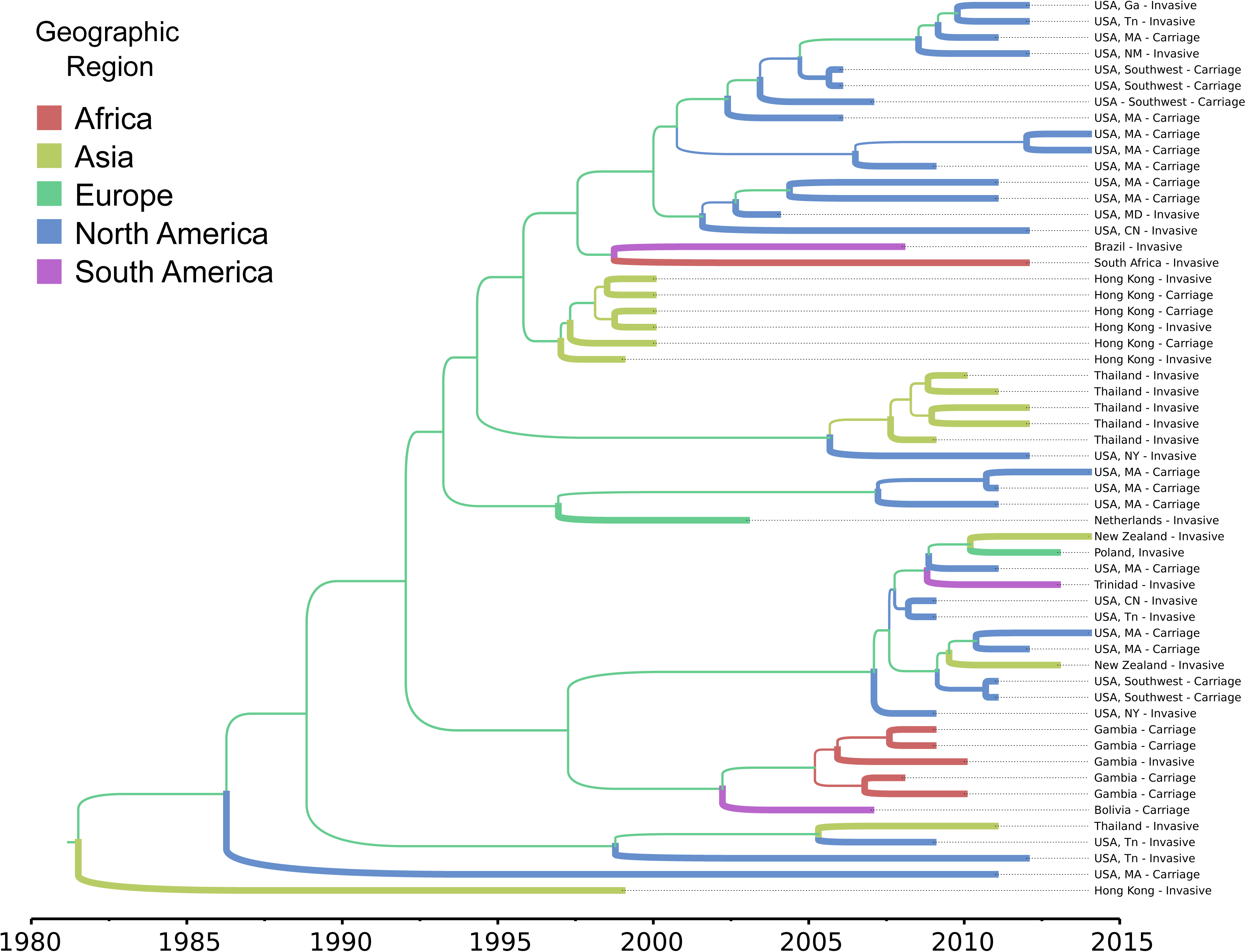
Maximum clade credibility, time-scaled phylogeny of CC180 serotype 3 (PMEN31) clade II isolates estimated using the structured coalescent model implemented in BEAST2. Branches are colored by geographic region and thickness is scaled values of posterior probabilities for geographic migration. Posterior probabilities for internal branches were all <0.30, precluding assessment of ancestral geographic dispersion.

**Figure 5.**
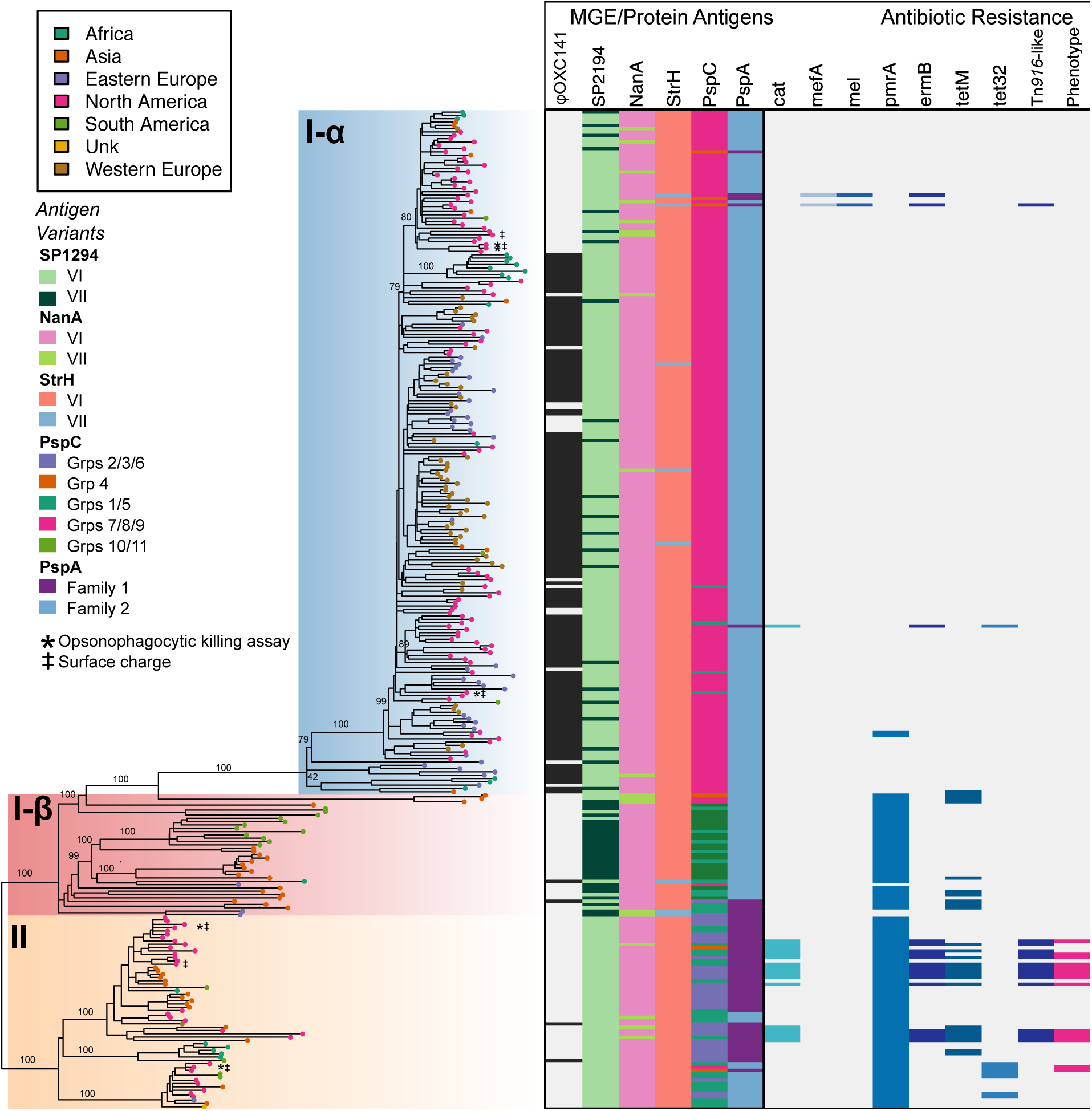
Phylogeny, polymorphic protein antigen variants, and antibiotic resistance. Midpoint rooted maximum-likelihood phylogeny corresponding to Figure 2A with geographic region of isolation represented as colored tip shapes. Clade I-α, I-β, and II are shaded consistent with Figure 2A. Bootstrap values from 100 replicates are provided for major clades and sub-clades (See supplemental phylogeny for all values). Strains used in opsonophagocytic killing assay (asterisk) and surface charge experiments (double dagger) are indicated on the phylogeny. Corresponding protein antigen variants for SP1294, NanA, StrH, PspC, and PspA are illustrated on the left half of the heatmap. Eight other protein antigens are excluded due to lack of variation in the sample. The presence and absence of AMR-associated genes are illustrated on the right half of the heatmap. The last column of the heatmap indicates genotypic antibiotic resistance that was confirmed by phenotypic testing (broth dilution or disk diffusion).

Clade II was found in 11 of 24 countries, 10 of which introduced PCV at some time during our study period (Figure 6). There was no association between countries that introduced PCV13 and the observation of an isolate belonging to Clade II (Fisher’s Exact, 4.2 95% CI: 0.3 – 239.9, p=0.3). There was also no correlation between year of vaccine implementation and the year of first identification of an isolate belonging to Clade II (Pearson’s correlation = 0.17, 95% CI: −0.52 – 0.72, p=0.64). Further, there was no clear pattern in vaccine introduction and Clade II presence.

**Figure 6.**
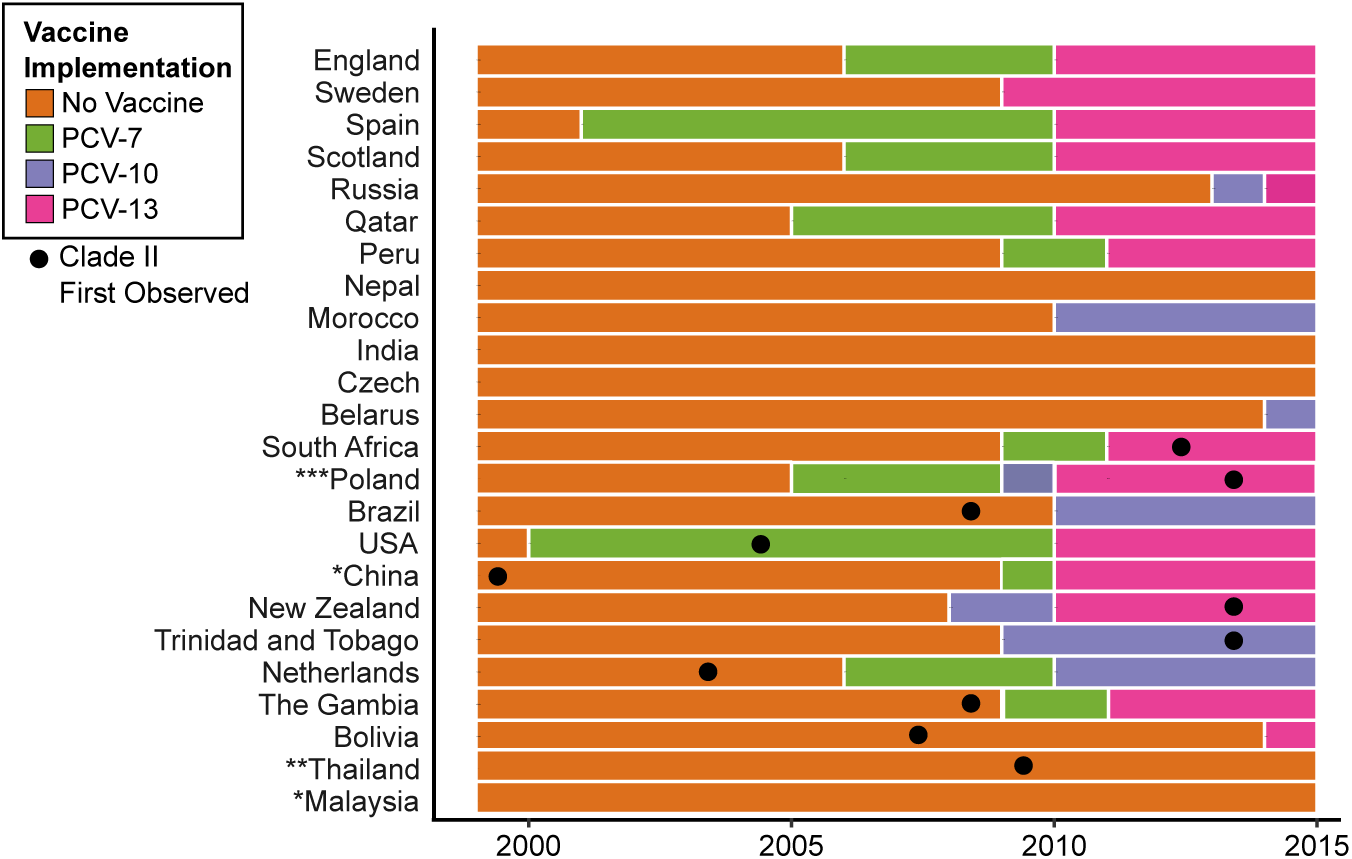
PCV implementation by county and year based on IVAC data from 1999–2014. For each country, the year is shaded based on PCV-7, PCV-10, and PCV-13 vaccine usage. Boxes marked with a dot designate the year in which a clade II isolate was first observed. *Malaysia and China have yet to introduce PCV as part of their national immunization program. Pneumococcal vaccine use in China varies regionally. Data from China reflects PCV use in Hong Kong where serotype 3 isolates were sampled. **PCV has been available in Thailand as an optional vaccine through the National Vaccine Program since 2006. However, uptake is <5% in children under 5. ***In Poland, PCV was available to parents through private pay. Population based vaccination, without catch-up campaign, was introduced in 2017 using PCV10.

### Recombination and genomic variation

We sought to identify whether genomic variation generated by recombination was associated with the recent increase of Clade II. We found that Clades I-α, I-β, and II experienced substantial recombination, impacting 1,179 CDS and diversifying gene content among clades (Supplementary Figures 5 and 6). The ratio of polymorphisms introduced through recombination compared to those introduced by mutation (*r/m*) for the entire sample was estimated at 1.76 with a mean recombination tract length of 12.3 kb. When independently assessed, we found Clade I-α possessed the lowest *r/m* among the three tested clades as well as the smallest tract length for recombination events (Table 2). Forty-two unique recombination events affecting 582 CDS occurred on the branch segregating clade II and I-β resulting in an *r/m* of 4.0 (Supplementary Figure 5). Comparison of *r/m* values between internal and terminal branches of the phylogeny suggests that the great majority of recombination events in CC180 are ancestral, although more recent events have occurred especially in Clade I-β (Table 2). We subsequently examined the *comCDE* operon, encoding the competence stimulating peptide and two-component regulatory system, and associated regulatory genes *comAB* to determine whether variations in recombination rates between Clades I-α, I-β, and II were associated with recombination or mutation among these loci. Overall, we found little diversity in the *comCDE* operon (mean pairwise SNP distance = 1 SE 0.5); however, *comD* possessed a non-synonymous mutation (AA104: Pro->Ser) among Clade I-β and II strains, which segregated them from Clade I-α. Further, while competence factor transporting protein *comA* was identical among the three clades, competence factor transport accessory protein *comB* was significantly diverged between Clades I-β/II and Clade I-α (mean pairwise SNP distance = 1.7 SE 0.7), suggesting phenotypic variation may have resulted from mutation or recombination. Assessment of recombination events identified that *comB* was located within a recombination event that affected all strains belonging to Clade II.

**Table 2.** Recombination among CC180 lineages. The number of polymorphisms introduced via recombination to mutation (*r/m*) and the number of recombination events to polymorphisms introduced by mutation (*ρ*/Θ) are reported for internal and terminal branches of dominant clades. Recombination occurring on internal braches is considered ancestral while terminal braches represent recent events. The overall recombination rates are also reported.

| Clade/s | Mean tract length (bp) | $r/m$ | | | $\rho/\theta$ | | |
| --- | --- | --- | --- | --- | --- | --- | --- |
|  |  | Internal | Terminal | Overall | Internal | Terminal | Overall |
| Clade I-α | 2,590.8 | 0.11 | 0.07 | **0.09** | 0.00 | 0.01 | **0.01** |
| Clade I-β | 11,457.6 | 4.15 | 2.33 | **3.35** | 0.03 | 0.02 | **0.03** |
| Clades I-β+II | 10,629.1 | 6.18 | 1.43 | **4.00** | 0.05 | 0.02 | **0.04** |
| Clade II | 9,446.5 | 5.34 | 0.32 | **2.67** | 0.05 | 0.01 | **0.03** |
| CC180 | 10,219.9 | 3.26 | 0.54 | **1.76** | 0.03 | 0.01 | **0.02** |

Recombination impacted a substantial proportion of the genome, focused in known recombination “hotspots” (Figure 7). As a result, we observed significant variation in mobile genetic elements (MGE) and gene content, including polymorphic protein antigens, evident through comparison of *de novo* assemblies and patterns of recombination. Pangenome comparison identified 1,437 core genes as well as variation in gene content among clades (Supplemental Figure 7). Two notable differences in MGE that resulted in gene content variation among CC180 clades were the presence of a Tn*916*-like conjugative transposon in Clade II strains (discussed below) and the absence of the 33.3 kb prophage OXC141 in Clades I-β and II [46,47]. The OXC141 prophage, which was putatively acquired by an ancestor of Clade I-α, has been lost multiple times by members of Clade I-α (Figure 5). Notably, it is absent from some Clade I-α isolates from North America, which includes a number of strains from Massachusetts that form a distinct subclade. In all, OXC141 is present in 71% of Clade I-α isolates and 50% of CC180 serotype 3 strains overall.

**Figure 7.**
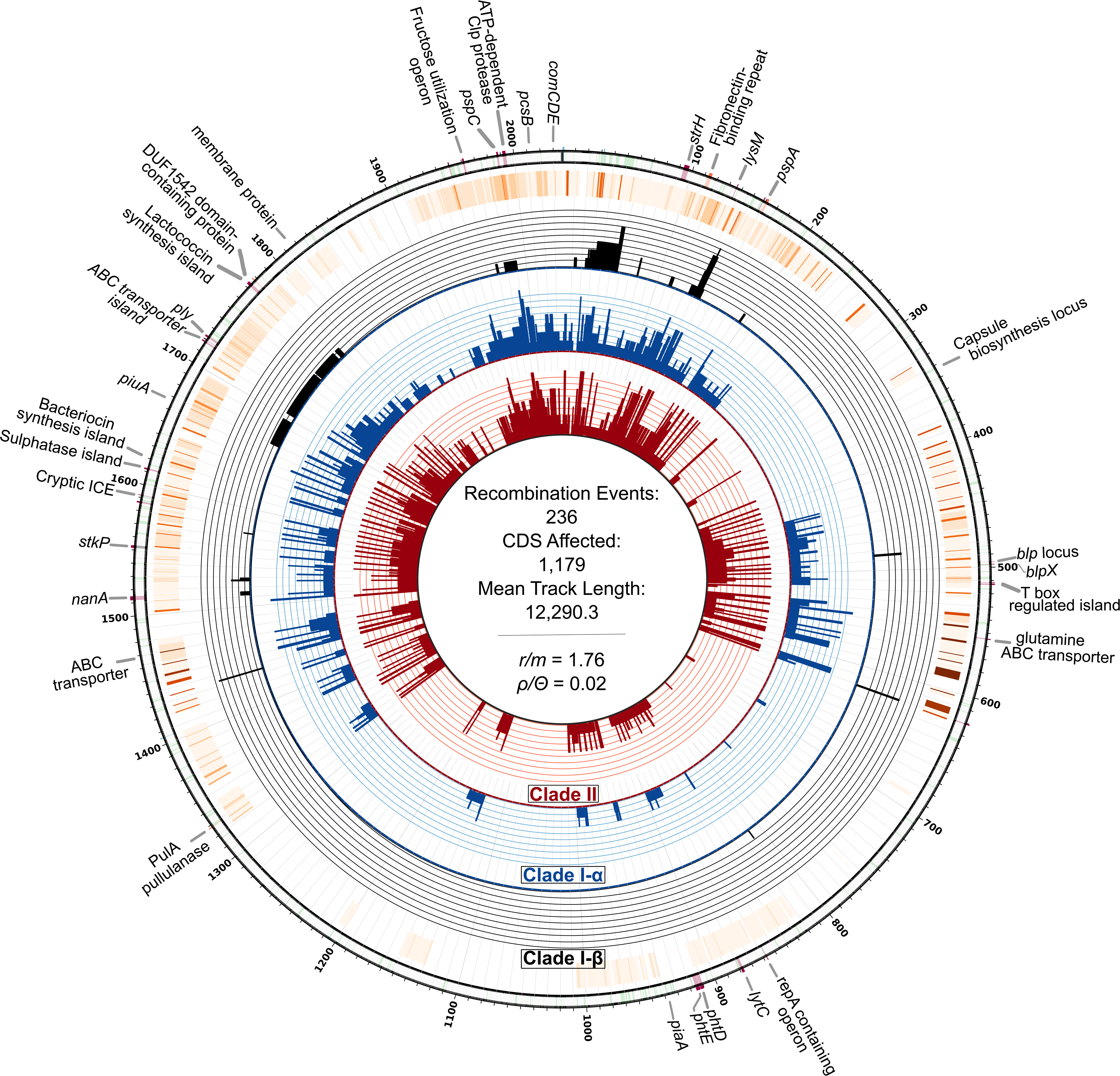
Circos plot of recent and ancestral inferred events inferred among CC180 serotype 3 isolates. Moving from the inner ring outward, rings show a histogram of unique recombination events occurring among isolates belonging to Clades I-β, I-α, and II, respectively, followed by a heatmap of cumulative recombination events. Finally, the outermost ring displays annotations of notable genes and genomic regions corresponding to the location on the OXC141 reference genome.

Among 13 tested polymorphic protein antigens, five were found to be variable, including membrane associated protein SP2194, cell-wall anchored proteins neuraminidase A (*NanA*) and β-N-acetylhexosaminidase (*StrH*), and surface exposed proteins *pspC* and *pspA* (Figure 5). SP2194 encodes the ATP-binding subunit of Clp protease and is involved in the expression of the *comCDE* operon, which plays an important role in competence, survival, and virulence of pneumococci [48]. Three internal and three terminal branches contain recombination events that have impacted SP2194. Major events occurred on each internal branch leading to Clades I-α, I-β, and II, and as a result, SP2194 is significantly diverged among the three clades (mean between clade SNP difference = 31.1 SE 3.8; p-distance 0.013 SE 0.002). Protein antigen genes *pspA* and *pspC* are known recombination “hotspots” [49], and here we find that recombination generated variation that subsequently become fixed in dominant clades and sub-clades. Family 2 *pspA* variant was largely associated with Clades I-α and I-β, while the Family I variant was associated with Clade II [50]. Using the *pspC* groupings based on structural and nucleotide variation proposed by Iannelli *et al.*, we found that Clades I and II had non-overlapping sets of *pspC* variation: Clade I possessed a *pspC* variant consistent with Groups 7/8/9, whereas Clade II variants include Groups 1/5 and 2/3/6 [51]. Hence, we have good evidence that the clades present divergent protein antigenic profiles to the human immune system.

### Antimicrobial-resistance (AMR) associated genes

We found that variations in AMR-associated genes contributed to differences in genome content among clades. Clade II isolates from Hong Kong (n=6) and USA (n=9) harbored a 37.6 kb Tn*916*-like conjugative transposon possessing *tetM* and *ermB* (Supplemental Figure 8). These strains also possess chloramphenicol acetyltransferase (*cat*). For 13 of 15 strains with Tn*916*, antibiotic susceptibility testing was available in GPS metadata or from previous publications [15]. In general, Tn*916* positive isolates exhibited high-level resistance to tetracycline, clindamycin, and erythromycin. However, three USA strains were susceptible to macrolides as the result of a previously described *ermBS* marker, a missense substitution within *ermB* that is associated with erythromycin and clindamycin-susceptibility [52]. Additional phylogenetic analysis of 26 core transposon clusters of orthologous genes (COGs) as well as genealogies of *Int-Tn, ermB*, and *tetM* illustrated that there were in fact two acquisitions of genetically similar Tn*916* transposons among Clade II isolates (Supplemental Figure 8). In both of these instances, the transposon was first observed in a Clade II isolate collected from Hong Kong, which lie basal to a clade of USA isolates possessing the respective variant.

Other notable differences between clades include the presence of *tet32* in eight Clade II isolates, which is distinct from *tetM* carried on Tn*916* transposons. In addition, the multi-drug efflux-pump *pmrA* was present in almost all Clade I-β and II isolates. We were able to confidently type PBP transpeptidase domains of 270 isolates to predict penicillin resistance. All were predicted to be penicillin-susceptible; however, four Clade II isolates possessed a PBP profile with first-step β-lactam resistance (transpeptidase domain profile 2–0-111, predicted MIC of 0.06). These four isolates also possessed Tn*916*. PBP profiles further differentiated CC180 clades, with the majority of Clade I-α isolates possessing profile 2–3-2, and Clades I-β and II strains profile 2–0-2 (Supplemental Table 2).

### Capsular Polysaccharide Variation

Interrogation of inferred recombination events showed that the CPS loci were unaffected by homologous recombination (Figure 7). Furthermore, while the ML phylogeny estimated from the alignment of CPS loci was clearly segregated between the dominant clades, the average pairwise nucleotide difference was only 1.6 (SE 0.4) SNPs (Supplemental Figure 9). Having ruled out significant genotypic variation, we continued to assess phenotype. While the surface charge of clade II isolates were slightly more negative among tested strains this difference was not significant (Supplemental Figure 10A). In addition, comparison of isolates from Clades I-α and II also found no significant difference in capsule shedding (Supplemental Figure 10B). Last, we observed neutrophil-mediated opsonophagocytic killing of Clade II isolates in the presence of antisera against both PCV13 and type 3 polysaccharide. In contrast, Clade I-α isolates were not susceptible to killing in the presence of type 3 polysaccharide antisera, and only one out of the three tested Clade I-α isolate was killed in the presence of high titer PCV13 antisera.

## Discussion

Serotype 3 pneumococci belonging to CC180, the most dominant serotype 3 clone globally, remain epidemiologically important. Over the last decade, Clade I-α and Clade II have increased in effective population size, with Clade II becoming particularly prevalent in Asia and North America and emerging in other regions where it has historically not been observed. Phylogenetic analysis demonstrates that Clade II is significantly diverged in both core genome nucleotide diversity and genome content, with multiple recombination events having occurred at the base of the lineage. Temporospatial surveillance data, coalescent analysis, and molecular epidemiology evidence suggests that the diaspora of Clade II may have originated in Southeast Asia in the 1990s and subsequently spread worldwide over two decades, concomitantly growing in prevalence. In North America, Clade II now makes up more than half of the post-PCV13 serotype 3 CC180 population. While neither use of PCV at a country level, nor timing on PCV introduction appears to be clearly associated with the rise of Clade II, we find that Clade II has divergent surface protein antigens and increased prevalence of antibiotic resistance compared to the other CC180 clades. These two observations are likely contributors to the recent increase in prevalence of Clade II.

While phylogeographic analysis was unable to confidently infer the ancestral migration events that disseminated Clade II, the presence of Tn*916* among strains from Hong Kong and the United States provides molecular evidence of migration from Asia to North America. In at least two instances, distinct Tn*916* transposons are found among isolates from Hong Kong, which are basal to strains from North America. This suggests the acquisition of this transposon predates the emergence of Clade II in North America and that strains circulating in Asia were the ancestral population of a large proportion (50%) of North American Clade II isolates. Further, multiple introductions of serotype 3 Clade II isolates to the USA from Asia may have occurred. Tn*916* confers macrolide resistance, with the rare exception of strains possessing a mutation in *ermB* [53]. Macrolide resistance has previously been described among serotype 3 ST180 isolates from Japan and Italy in 2003 and 2001, respectively, where macrolide-resistant serotype 3 strains are prevalent [13,54,55]. In addition, macrolide and tetracycline resistance ST180 strains have been reported from Taiwan (1997), Spain (2002), and more recently, Canada (2011–14), and Germany (2012) (https://pubmlst.org/). Unfortunately, MLST is not discriminating enough to distinguish between the CC180 lineages we discuss here, and as genomic data are not available for these isolates, we cannot determine how they relate to our sample. However, as macrolide resistance among CC180 serotype 3 isolates is relatively rare and largely isolated to Clade II, the early identification of macrolide resistance strains from Hong Kong, Taiwan, and Japan may represent the ancestral population of Clade II putatively possessing Tn*916*, further supporting a Southeast Asian origin and implicating antibiotic resistance as a possible contributor to the recent success of Clade II.

As PCV use has previously been associated with major changes in pneumococcal population structure [56,57], we considered that serotype 3 may have increased in those settings where PCV was introduced. Further, variation in effectiveness of the serotype 3 component of PCV13 among CC180 clades may have disproportionally precipitated an increase in Clade II. Here, we find no evidence that PCV use, albeit at a country level, has shaped the serotype 3 CC180 population. However, a limitation is that these comparisons ignore important factors such as phylogenetic population structure (i.e., whether Clade II was already present), age of cases, and details of vaccine roll-out (e.g., timing, targeted age groups, and vaccine coverage). Unfortunately, these data are incomplete for a number of cases/countries, precluding an depth analysis of the putative association of PCV with emergence of Clade II. In absence of a clear epidemiological etiology for the increase of Clade II, we investigated phenotypic and genomic variation between Clades I-α and II.

Historically, antibiotic resistance among serotype 3 CC180 strains has been low. This is because the previous population was dominated by isolates from Clade I, which is largely devoid of AMR-associated genes and mutations. This may result from the high invasiveness of serotype 3 and its low carriage duration, which in turn reduces antibiotic exposure [58]. Here we find chloramphenicol, macrolide, tetracycline, and first step penicillin resistance more frequent among Clade II isolates, providing one possible explanation for its recent increase in prevalence and emergence in previously unobserved regions. In fact, increased macrolide usage in North America has previously been associated with an increase in macrolide resistant *Staphylococcus aureus* and nonsusceptible *S. pneumoniae* [59,60].

In addition to variations in AMR-associated genes, we also find that MGE and polymorphic protein antigens varied among Clades I-α, I-β, and II. The previously described OXC141 was generally absent among Clades I-β and II and has been lost by a North American sub-clade of Clade I-α [46,47]. Temperate bacteriophages may be associated with fitness defects among pneumococci or changes in virulence and competence [61,62]. Whether the presence or absence of OXC141 relates to the relative success of serotype 3 CC180 clades is unclear. However, the absence of OXC141 from Clade II as well as the loss of the phage by a subclade of North American Clade I-α strains remains notable and warrants further investigation. Contributing to genomic variations among clades, *pspA, pspC,* and SP2194 (ATP-dependent Clp protease) were found to vary. Natural immunity to protein antigens is generated through nasopharyngeal colonization, and this immunity may protect against colonization and invasive disease [22,63]. Protein antigen variants were generally conserved among Clade I-α isolates and varied in Clades I-β and II. As protein antigen variants are serologically distinct, variations in antigen profiles among CC180 clades could impact relative transmissibility and virulence, resulting in differences in carriage duration, transmission, and invasive capacity [22,51,64,65]. For example, in previous comparisons of Family 1 and 2 *pspA* variants in isogenic strains of serotype 3 (WU2), Family 2 mutants were slightly less virulent and bound less human lactoferrin, impacting nasopharyngeal colonization [66]. Here, we find that Clade II isolates largely possess the Family 1 *pspA* variant. Further, variants of *pspC* differentially bind factor H and generate distinct antibody responses, promoting immune escape [67]. Most isolates in Clade I-α possess a *pspC* variant corresponding to Groups 7, 8, and 9, which have been grouped here because they share the same structural organization [51]. In contrast, Clades I-β and II possess multiple polymorphic *pspC* variants interspersed throughout the topology of the clades, suggesting that multiple recent recombination events have generated variation in this known recombination hotspot as well as a diversified antigenic profile. Taken together, differences in antigenic profiles add to the phenotypic variability among CC180 clades, which may lead to increased carriage duration and immune escape of Clade II isolates. While the epidemiological implications of these finding require further investigation, together with antibiotic resistance, antigenic variation presents another possible contributor to the success of Clade II.

The population of serotype 3 has been previously thought to be largely clonal, clustering into a group of closely related PFGE patterns corresponding to the Netherlands^3^–31 clone. Further, evolution of CC180 was thought to be driven primarily through nucleotide substitution and estimates of recombination rates based largely on Clade I-α suggested a comparatively lower rate to other PMEN clones [12], mainly dominated by micro-recombination events [68]. However, the expanded global sample of PMEN31 serotype 3 genomes demonstrates in detail how the prevalent lineages have diverged following significant ancestral recombination events. While comparatively rare, recombination continues to shape the CC180 population, as terminal branches in all three clades show evidence of recent events. This inferred clonality likely resulted from the limited sample of type 3 isolates from restricted geographies. Further, while serotype 3 is generally understood to show lower competence than other serotypes based on *in vitro* studies [12,69], we found that the emergent Clades II and I-β have more evidence of historical recombination events (ρ/Θ) and a greater proportion of macro-recombination events than the previously dominant Clade I-α. Therefore, Clade I-α may possess a genomic variant that reduces its competence, a mutation that is absent in Clade I-β and Clade II. Consistent with this hypothesis, the Clade I-α *comCDE* operon generates an antisense transcript, an observation that likely relates to the low observed recombination rate of Clade I-α [12]. We show amino acid changes in the *comCDE*, specifically *comD*, segregate Clade I-α from I-β, and II. We also found variation in the Clp protease gene, which regulates the *comCDE* operon, and is therefore associated with competence rates among pneumococci and other bacteria [48,70]. We therefore suggest that variation in the competence machinery of Clade II likely contributes to increased recombination rates, and this has generated a more diverse antigenic profile among Clade II isolates and allowed them to acquire multi-drug resistance [71,72]. Indeed, higher rates of recombination have been associated with antimicrobial resistance [73].

Because serotype 3 pneumococcal strains possess a mucoid capsule and release significantly more capsular polysaccharide during *in vitro* growth and infection in mice, it is hypothesized that this soluble polysaccharide absorbs anti-capsular antibody, effectively impeding antibody-mediated killing *in vitro* and *in vivo* [18,20,45]. We found that Clades I-α and II both shed CPS during growth; however, the amount of CPS released did not significantly differ between the two clades. Another phenotype that has been found to correlate with increased survival due to resistance from phagocytosis by human neutrophils, and contribute to success in nasopharyngeal carriage is low surface charge, measured as zeta potential [19]. We therefore compared this between Clades I-α and II, finding no significant difference in charge between clades, suggesting capsular properties are similar. Where we did find a significant phenotypic differences between the clades, it was not easy to relate to the apparent increase in Clade II following PCV13 introduction; we initially hypothesized that antisera against PCV13 and type 3 polysaccharide might be more effective against isolates from Clade I-α, explaining its apparent decline. For both Clades I-α and II, neutrophil-mediated opsonophagocytic killing occurred only at high antibody levels. Clade I-α isolates appeared to be resistant to killing by PCV13 antisera, while Clade II isolates were more readily killed. The resistance of Clade I isolates to opsonophagocytic killing is consistent with the low efficacy of PCV13 on serotype 3 invasive disease incidence [9,74], but difficult to link to the recent emergence of Clade II. However, these results should not be considered a direct representation of vaccine effectiveness, because opsonophagocytic killing may be a poor proxy for the ability to colonize and transmit, and moreover killing only occurred at high titers. Our findings are consistent with those of other groups, suggesting that higher antibody concentrations are needed for killing *in vitro* [45]. They also demonstrate that closely related bacteria with the same capsule do not necessarily behave similarly in the face of vaccination.

Together, our findings further support the involvement of anti-protein antibodies in mediating anti-serotype 3 immunity. Anti-protein antibodies may act in synergy *in vivo* with anticapsular antibodies to mediate pneumococcal killing. This is an intriguing hypothesis in light of our other findings about the differences in protein variant composition among serotype 3 strains. These differences in protein composition underscore the need for better evaluation of the role anti-protein antibodies in mediating anti-pneumococcal protection. Overall, the role of these in recent dynamics of serotype 3 epidemiology requires further study.

Serotype 3 remains an epidemiologically important serotype, continuing to cause invasive disease and increasing in carriage prevalence in North America. Here, we explore multiple epidemiological, genotypic, and phenotypic factors associated with the persistence of serotype 3 in carriage and disease. We found that the recent success of CC180 is related to an increasing clade, which is distinct from the previous population in antigenic composition, antibiotic susceptibility, and competence. Based on OPKA, we also find evidence to support the low observed efficacy of PCV13 against serotype 3, which was previously dominated by CC180 Clade I-α in Europe and North America. Optimistically, the efficacy of PCV13 against the emergent Clade II may be higher; however, more *in vitro* and *in vivo* studies are required to confirm this finding.

Our study is not without limitation. While we base our analysis on the largest collection of extant CC180 genomes, geographic and temporal sampling variations exist. To mitigate coalescent analysis bias, we down-sampled according to published approaches. Additionally, while the clonality and low diversity of Clade I-α may, at first approximation, suggest a more recent origin, HPDs for the TMRCA of Clade I-α and II estimated through coalescent analysis are non-overlapping and correspond to previously published estimates [12]. Further, despite modest sampling from multiple geographies prior to 1999 (n=34), Clade II isolates were not observed. Our results instead indicate that the higher recombination rate and recent population expansion of Clade II has led to its greater diversity. Last, our genomic results may be affected by the accuracy of bioinformatic programs. Wherever possible, results were verified using multiple approaches or manual inspection of the data.

Overall, our analysis emphasizes the need for routine, representative sampling of isolates from disperse geographic regions, including historically under-sampled areas. We also highlight the value of genomics in resolving antigenic and epidemiological variations within a serotype, which may have implications for future vaccine development. As PCV usage and antibiotic consumption expands globally, it is imperative to continue genomic and epidemiological surveillance of pneumococci to detect the emergence of new lineages and monitor changes in clinical presentation and severity, as these data have direct implications for prevention and management of pneumococcal disease.

## Acknowledgements

We acknowledge the Wellcome Trust Sanger Institute sequencing facility. We would also like to thank Richard Malley’s Laboratory for providing the antisera for the OPKA assays.

**Supporting Information Legends**

S1 – Supplemental methods, tables (2), and figures (11)

S2 – File containing accession numbers and associated metadata

